# Delineation of a molecularly distinct terminally differentiated memory CD8 T cell population

**DOI:** 10.1101/2020.07.17.207266

**Authors:** J. Justin Milner, Hongtuyet Nguyen, Kyla Omilusik, Miguel Reina-Campos, Matthew Tsai, Clara Toma, Arnaud D. Delpoux, Brigid S. Boland, Stephen M. Hedrick, John T. Chang, Ananda W. Goldrath

## Abstract

Memory CD8 T cells provide durable protection against diverse intracellular pathogens and can be broadly segregated into distinct circulating and tissue-resident populations. Paradigmatic studies have demonstrated circulating memory cells can be further divided into effector memory (Tem) and central memory (Tcm) populations based on discrete functional characteristics. Following resolution of infection, we identified a persisting antigen-specific CD8 T cell population that was simultaneously terminally-fated with potent effector function but maintained memory T cell qualities and conferred robust protection against reinfection. Notably, this terminally-differentiated effector memory CD8 T cell population (terminal-Tem) was conflated within the conventional Tem population, prompting redefinition of the classical characteristics of Tem cells. Murine terminal-Tem were transcriptionally, functionally, and developmentally unique compared to Tem cells. Through mass cytometry and single-cell RNAseq analyses of human peripheral blood from healthy individuals, we also identified an analogous terminal-Tem population of CD8 T cells that was transcriptionally distinct from Tem and Tcm. A key finding of this study was that parsing of terminal-Tem from conventionally defined Tem challenges classical characteristics of Tem biology, including enhanced presence in lymphoid tissues, robust IL-2 production and recall potential, greater than expected homeostatic fitness, refined transcription factor dependencies, and a distinct molecular phenotype. Classification of terminal-Tem and clarification of Tem biology hold broad implications for understanding the molecular regulation of memory cell states and harnessing immunological memory to improve immunotherapies.

## Introduction

Memory CD8 T cells are critical mediators of long-lived immunity and provide dynamic protection against intracellular pathogens and malignancies. Given the unique attributes of memory T cells, including specificity, durability, and rapid effector functions, leveraging this population of cells is a key objective of diverse immunotherapies and vaccines. Memory CD8 T cells can be broadly segregated into recirculating Tcm and Tem subsets predominantly found in the blood and lymphoid tissues as well as tissue-resident memory T cells (Trm), which are primarily localized to non-lymphoid sites (1). Numerous studies have established that Tcm and Tem populations are phenotypically and functionally distinct. Tcm are generally considered to exhibit greater homeostatic proliferation and longevity, multipotency, recall potential, and lymph node homing capacity due to characteristically elevated expression of CD62L and CCR7 (1–6). Conversely, Tem are thought to be relatively shorter-lived and terminally-fated, exhibit an effector phenotype marked by granzyme production, and primarily localized in the vasculature relative to lymphoid tissues and in certain contexts to recirculate through or populate non-lymphoid sites (2, 7–9). This partitioning of functional attributes and localization properties within the circulating memory compartment provides a division of labor as well as flexibility in the mounting of robust defenses to diverse pathogens and neoplasms.

Memory T cells are primarily derived from a multipotent memory precursor (MP) population distinguished by elevated expression of the IL-7 receptor (CD127) and low expression levels of KLRG1, whereas CD127^lo^KLRG1^hi^ terminally-differentiated effector (TE) cells are generally considered to be short-lived and terminally-fated following acute infection (10–12), although ex-KLRG1-expressing cells have been shown to form multiple memory populations (13). As our understanding of T cell states and fates has expanded, it has become evident that diverse intracellular and extracellular cues dictate CD8 T cell differentiation and homeostasis, ultimately through the dynamic activity of fate-specifying transcription factors (3, 14). Canonical pro-effector transcription factors linked to the formation of TE cells and Tem include T-bet (11), Blimp1 (15, 16), Zeb2 (17, 18), Stat4 (19), and Id2 (20–23), whereas pro-memory transcription factors required for optimal MP and Tcm differentiation include Eomes (24), Bcl6 (25, 26), Foxo1 (27–29), Stat3 (26), and Id3 (22, 30). However, our ability to probe the molecular underpinnings of Tcm and Tem fates is limited by our capacity to accurately define and delineate these distinct memory populations.

In humans, Tem and Tcm are frequently defined as CD45RO^hi^CCR7^lo^ and CD45RO^hi^CCR7^hi^, respectively (1, 4, 5, 31). However, our understanding of the function, differentiation, and molecular regulation of memory T cells is primarily derived from mouse models, wherein a common strategy for discriminating Tem and Tcm is often based solely on CD62L expression (4, 8, 32–40). A number of studies have reported heterogeneity within the Tem compartment, including a persisting “effector-like” population of KLRG1^hi^ cells (41), CD43^lo^CD27^lo^ cells (42), or CX3CR1^int^ peripheral memory cells (43), and some reports suggest alternative or complimentary (in addition to CD62L for example) approaches for demarcation of memory populations based on differing levels of CX3CR1 (43, 44), CXCR3 (43, 45), CD27 (4, 46), CD28 (46), KLRG1 (13, 17), and CD127 (10, 47, 48). Here, we report that the conventional CD62L^lo^ Tem population is conflated with a transcriptionally and functionally distinct population of CD127^lo^CD62L^lo^ terminally differentiated effector memory cells (terminal-Tem). This finding has a number of implications, but the principal relevance is that the general understanding of the phenotype, function, and transcriptional regulation of Tem cells has been confounded by a contaminating terminal-Tem population, requiring re-examination of Tem biology. Further, we show terminal-Tem exhibit potent cytotoxic activity but have limited multipotency and recall potential. Thus, we here argue for a definition of Tem as CD127^hi^CD62L^lo^ and provide a framework for ascribing novel (or previously muddled) characteristics of Tem including an enhanced presence in lymphoid tissues, robust IL-2 production and recall potential, greater than expected homeostatic fitness, and a distinct molecular phenotype. Additionally, we refine canonical roles for the key lineage-specifying transcription factors T-bet, Blimp1, Bcl6, and Foxo1. Last, we identify an analogous terminal-Tem subset within the human circulating memory CD8 T cell compartment with discrete transcriptional and phenotypic qualities compared to Tem, Tcm, and effector cells. Given that memory T cells are linked to prevention and progression of diverse diseases, an in-depth understanding and elucidation of circulating memory states hold widespread implications from molecular analyses to clinical studies and allow greater context for pinpointing which memory state might be most therapeutically relevant for a given disease setting.

## Results

### Terminal-Tem are a distinct subset of CD8 T cells contained within the conventional CD62L^lo^ Tem population

Tem are most often classified as CD62L^lo^CCR7^lo^ and Tcm as CD62L^hi^CCR7^hi^ (1, 3, 31). However, CCR7 staining is not typically incorporated in murine memory CD8 T cell subsetting as the staining process (43) is not as straightforward as for CD62L, and both molecules are generally co-expressed (4, 5). Therefore, a majority of the studies investigating circulating memory T cell populations utilize differential CD62L expression as the defining molecule to distinguish Tcm and Tem (4, 8, 32–40). Expression of CD127 has also been widely used to define antigen-experienced CD8 T cell populations with memory qualities (10, 16, 47–49). Utilizing the LCMV infection system, we found that LCMV GP_33-41_-specific tetramer^+^ memory CD8 T cells (*SI Appendix*, Fig. 1 *A*) or TCR transgenic P14 cells with low expression levels of CD62L (i.e. conventional Tem) were heterogeneous for CD127, memory-associated costimulatory molecule CD27, and the TE- or short-lived effector-associated molecule KLRG1 (Fig. 1 *A*). Additionally, circulating antigen-specific CD62L^lo^ memory cells generated following *Listeria monocytogenes* infection also displayed this heterogeneity (*SI Appendix,* Fig. 1 *B*). More than two months after LCMV infection, >30% of CD62L^lo^ cells remained CD127^lo^ and CD27^lo^ (Fig. 1 *A*) prompting the questions: Are CD127^lo^ cells at a ‘memory timepoint’ considered memory cells? Should the CD127^lo^ cells be grouped into the conventional ‘Tem’ population? How do these CD127^lo^ cells fit with other strategies for distinguishing memory T cell subsets (e.g. CX3CR1, CD43, CD27)?

**Figure 1.**
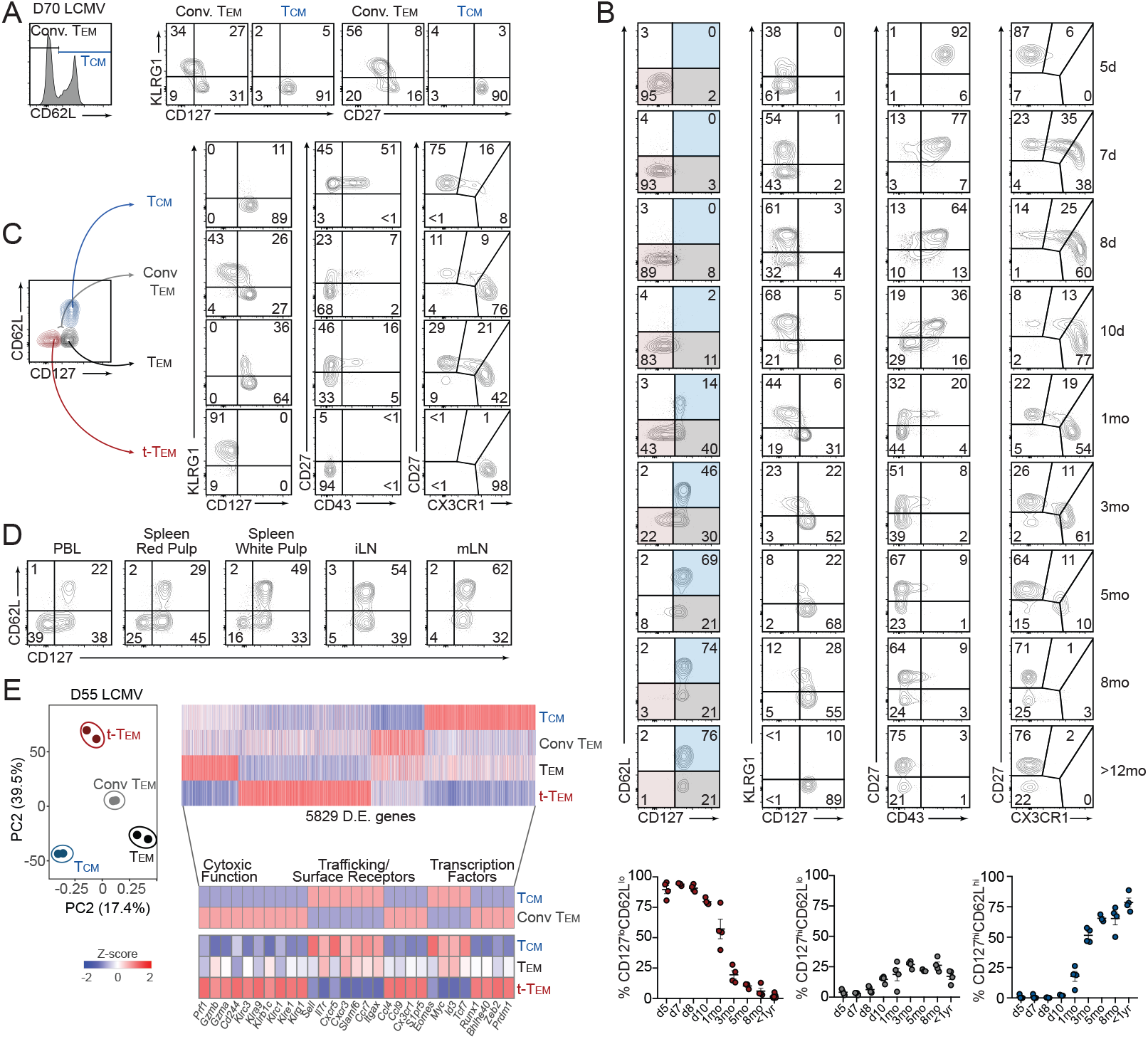
Terminal-Tem are a distinct subset of CD8 T cells contained within the conventional Tem population. P14 CD8 T cells were adoptively transferred into congenically distinct recipient mice that were subsequently infected with LCMV. (*A*) Expression of KLRG1, CD127 (left), and CD27 (right) by Tcm (CD62L^hi^) and conventional (Conv.) Tem (CD62L^lo^) within peripheral blood lymphocytes (PBL) on day 70 of infection. *(B)* Expression of CD62L, CD127, KLRG1, CD27, CD43, and CX3CR1 (top) by P14 cells in PBL at indicated times of LCMV infection. Frequency of CD127^lo^CD62L^lo^ (highlighted red), CD127^hi^CD62L^lo^ (highlighted grey), and CD127^hi^CD62L^hi^ (highlighted blue) in PBL following LCMV infection (bottom). *(C)* Representative expression patterns of KLRG1, CD127, CD27, CD43, and CX3CR1 by terminal-Tem, Tem, Conv. Tem, and Tcm. *(D)* Representative flow cytometry plots demonstrating the frequency of terminal-Tem, Tem, and Tcm in indicated tissues (>30 days post-infection). Splenic red and white pulp localized P14 cells were discriminated by intravascular staining of CD8α. *(E)* On day 55 of infection, splenic terminal-Tem, Tem, Conv. Tem, and Tcm P14 cells were sorted for RNA-seq analysis. Principal component analysis of gene expression from the sorted memory populations (left), and heatmap illustrating 6467 differentially expressed genes (≥1.5-fold) ordered through hierarchical clustering (top right) among terminal-Tem, Tem, Conv. Tem, and Tcm, or highlighted key genes (bottom). Numbers in plots are the frequency of cells in the indicated gate. All data are from 2 independent experiments with n=3-5 per timepoint, and RNAseq samples consist of 2 biological replicates wherein each replicate is comprised of cells pooled from 2 mice. Graphs indicate mean ± s.e.m, and symbols represent an individual mouse *(B)*.

In attempt to develop a comprehensive and unifying understanding of the dynamics of well-studied molecules often used to distinguish circulating memory CD8 T cell populations, we measured expression patterns of CD62L, CD127, KLRG1, CD43, CD27 and CX3CR1 over the course of >1 year following LCMV infection. Although CD43 and CD27 have previously been reported to distinguish circulating memory cells with differing levels of effector function and recent investigations have established varying CX3CR1 levels can delineate distinct populations of circulating memory cells, CD62L^hi^ Tcm and conventional CD62L^lo^ Tem classification is widely used to subset circulating memory populations. Thus, an in-depth characterization of how so-called ‘memory markers’ fully describe memory CD8 T cell states is needed. As expected, the proportion of CD127^hi^ circulating memory T cells increased over time after LCMV infection, and 1yr after infection nearly all P14 cells were CD127^hi^ in the blood (Fig. 1 *B*). We noted a prominent CD127^lo^CD62L^lo^ population of cells persisting up to 5 months after infection, and therefore sought to investigate this apparent heterogeneity within the CD62L^lo^ population of memory cells as well as to clarify the molecular identity, ontogeny, and function of Tem in relation to the CD127^lo^CD62L^lo^ population.

CD127^lo^CD62L^lo^ cells at memory timepoints (>30 days after infection) exhibited expression patterns consistent with a terminal effector phenotype, including elevated KLRG1 expression, low expression of CD43 and CD27, and robust expression of CX3CR1; furthermore, while longer lived than TE cells, the persisting CD127^lo^CD62L^lo^ population ultimate wanes after 3-5 months and thus have limited durability compared to CD127^hi^CD62L^lo^ and Tcm populations (Fig. 1 *B*). Therefore, we refer to CD127^lo^CD62L^lo^ cells as terminally-differentiated effector memory T cells (terminal-Tem, Fig. 1 *C*). We also redefine Tem as CD127^hi^CD62L^lo^ cells and Tcm as CD127^hi^CD62L^hi^, although CD62L staining alone is sufficient to distinguish Tcm, as nearly all CD62L^hi^ cells are CD127^hi^. We next utilized this revised memory T cell nomenclature to investigate the phenotypic characteristics and unique qualities of circulating memory T cells with an emphasis on contrasting terminal-Tem with Tem.

Following LCMV infection, terminal-Tem were primarily localized in the vasculature and less abundant in lymph nodes (Fig. 1 *D*), consistent with reports of CX3CR1^hi^ cells predominating in the blood (43). However, we find a considerable proportion of CD127^hi^CD62L^lo^ Tem localized not only in the blood but also in splenic white pulp and lymph nodes as well. Thus, clarifying conventional Tem into discrete CD127^hi^CD62L^lo^ Tem and CD127^lo^CD62L^lo^ terminal-Tem subsets reveals relevant biology about memory T cells; classically, conventional Tem abundance in lymphoid tissues was thought to be relatively lower than blood (2, 3), but here we demonstrate that this was chiefly due to a contaminating terminal-Tem population and that Tem in steady state conditions are abundant in the spleen and draining lymph nodes.

We next profiled the transcriptome of the newly defined terminal-Tem and CD127^hi^CD62L^lo^ Tem populations compared to Tcm and conventional CD62L^lo^ Tem to understand their transcriptional relationship as well as to elucidate differential gene-expression programs contributing to the fate and homeostasis of these populations. Principal component analysis illustrated that, indeed, terminal-Tem are transcriptionally distinct from Tcm and Tem, but importantly also demonstrated that redefined Tem and Tcm are distinct populations and not simply a uniform population of cells with varying expression levels of CD62L. We detected >5000 genes differentially expressed among the three memory subsets, and closer evaluation of mRNA expression levels of key functional and transcriptional molecules further confirmed that terminal-Tem displayed elevated expression of key cytotoxic, effector, and migratory molecules (Fig. 1 *E*). Inclusion of the transcriptional profile of conventionally defined Tem (CD62L^lo^) in the transcriptomic analyses highlighted how redefining Tem provides novel insight into regulatory gene-expression programs and the lineage relationship of memory T cells. For example, contrary to current paradigms, Tem express relatively low levels of *Prdm1* and *Zeb2* but elevated levels of *Tcf7* and *Id3* after parsing of CD127^lo^CD62L^lo^ terminal-Tem and CD127^hi^CD62L^lo^ Tem populations (Fig. 1 *E*). Last, as Trm are a distinct memory T cell population with robust effector features (50, 51), we compared the transcriptional relationship of terminal-Tem, Tem, Tcm, and intestinal Trm, which also revealed unique gene-expression patterns between each population and supported the underlying notion that terminal-Tem are a discrete CD8 T cell population (*SI Appendix,* Fig. 2 *A-B*).

**Figure 2.**
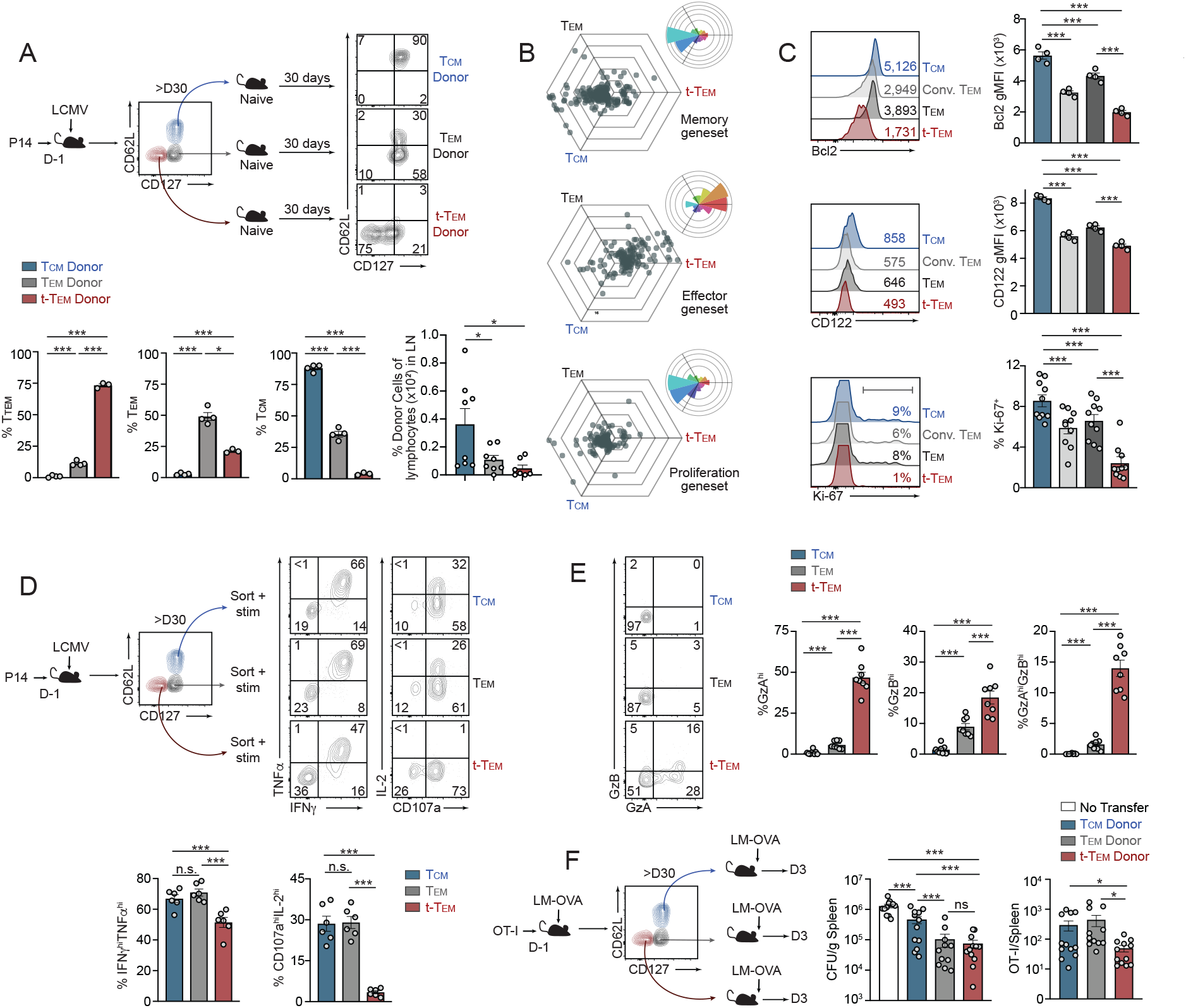
Distinct fate, homeostasis, and effector phenotype of CD127^lo^CD62L^lo^ terminal-Tem and CD127^hi^CD62L^lo^ Tem. *(A)* More than 30 days after LCMV infection, terminal-Tem (CD127^lo^CD62L^lo^), Tem (CD127^hi^CD62L^lo^), and Tcm (CD127^hi^CD62L^hi^) P14 memory subsets were sorted and transferred into naive, congenically distinct mice. After >30 days post-transfer, the frequency of each donor population in the spleen was analyzed by flow cytometry. Representative plots (top) and quantification of the phenotype of (bottom) is shown as well as frequency of donor cells recoevered from lymph nodes (bottom right). *(B)* Triwise comparisons of differentially expressed genes (≥1.5-fold) within the indicated gene-expression signature were plotted in a hexagonal diagram in which the magnitude of differential expression is reflected by distance from the origin. Rose plots (upper right corner of each hexagonal plot) indicate the percentages of genes in each orientation. Memory- (top), effector- (middle) and proliferation-associated (bottom) gene sets are displayed. *(C)* Flow cytometry analysis of terminal-Tem, Tem, Conv. Tem, and Tcm on day >D35 of infection. Representative plots (left) and quantification of gMFI or % positive (right) are shown. *(D)* Memory subsets were sorted and cultured *ex vivo* with GP_33-41_ peptide and congenically distinct splenocytes for assessment of TNFα, IFNγ, IL-2 production, and degranulation (CD107a). Representative flow cytometry plots (top) and quantification of the frequency of cytokine production by each subset (bottom) is shown. *(E)* GzA and GzB expression by terminal-Tem, Tem, and Tcm was analyzed by flow cytometry (left) and quantified (right). *(F)* OT-I CD8 T cells were transferred into congenically distinct mice that were infected with LM-OVA the following day. More than 30 days after infection, terminal-Tem (CD127^lo^CD62L^lo^), Tem (CD127^hi^CD62L^lo^), and Tcm (CD127^hi^CD62L^hi^) subsets were sorted and transferred to naive recipients that were challenged with LM-OVA. On day 3 of challenge, bacterial load in spleens was assessed (CFU/gram of tissue) and abundance of donor cells in recipient spleens were enumerated by flow cytometry. Numbers in plots represent gMFI or the frequency of cells in the indicated gate *(A, C-E).* All data are from one representative experiment of 3 independent experiments with n=3-4 per group *(A,C),* or data are combined from 3 independent experiments (Ki-67 staining in *C*, *F*), or data are combined from 2 independent experiments *(E)*, or combined from 3 independent experiments where each experiment consisted of 2 biological replicates and each replicate was sorted from 1 or 2 mice combined *(D)*; ***P<0.005. Graphs indicate mean ± s.e.m, and symbols represent an individual mouse or an individual replicate.

### Distinct fate, homeostasis, and effector phenotype of CD127^lo^CD62L^lo^ terminal-Tem and CD127^hi^CD62L^lo^ Tem

Profiling of population dynamics in the blood revealed that the terminal-Tem population declines over time while Tcm become the predominant memory population, as expected (Fig. 1 *B*). It was unclear if the loss of terminal-Tem was due to limited homeostatic proliferation, impaired survival, or trans-differentiation of terminal-Tem. Although we hypothesized limited flexibility in terminal-Tem fate given their resemblance to TE, we first tested if terminal-Tem were able to trans-differentiate into other memory populations, potentially contributing to the loss of terminal-Tem. Terminal-Tem, Tem, and Tcm were sorted and transferred into naive recipient mice, and approximately 30 days after adoptive transfer, the donor cells were phenotyped (Fig. 2 *A*). We found that terminal-Tem primarily retained a CD127^lo^CD62L^lo^ phenotype with minimal conversion to other memory subsets, consistent with their designation as a terminally-differentiated population. In contrast, approximately 50% of Tem were able to upregulate CD62L (ostensibly converting to Tcm), and transferred Tcm remained phenotypically stable, consistent with previous reports (4). The relative fixed fate of terminal-Tem was also emphasized by their limited presence in lymph nodes upon retransfer into naive recipient mice (Fig. 2 *A*).

As terminal-Tem exhibit relatively minimal trans-differentiation into other memory T cell populations, we next assessed their capacity for homeostatic proliferation and long-term survival. Through three-way comparisons, we found that terminal-Tem expressed lower levels of signature T cell proliferation genes (52) as well as memory signature genes, but elevated levels of characteristic terminal effector genes relative to Tem and Tcm (Fig. 2 *B*). Flow cytometry analyses confirmed the distinct transcriptional differences observed, further demonstrating terminal-Tem express relatively low levels of pro-survival molecules CD122 and Bcl2 and undergo lower levels of homeostatic proliferation indicated by diminished Ki-67 staining (Fig. 2 *C*). Taken together, the loss of the terminal-Tem population over time (Fig. 1 *B*) is likely due to diminished homeostatic fitness (i.e. the capacity for homeostatic proliferation and long-term survival).

We next sought to determine if terminal-Tem and the redefined Tem population are functionally distinct from each other as well as Tcm. To assay cytokine production, terminal-Tem, Tem, and Tcm populations were sorted (due to diminished surface abundance of CD127 and CD62L upon *ex vivo* stimulation (53)) and incubated with cognate peptide. Terminal-Tem exhibited reduced polyfunctionality compared to Tem and Tcm, with a lower frequency of IFNγ^hi^TNFα^hi^ cells (Fig. 2 *D*). Consistent with an impaired homeostasis phenotype, the percentage of IL-2 producing terminal-Tem was approximately 8-fold lower compared to Tcm and Tem, while in contrast to current paradigms (2) an equivalent frequency of Tem produced IL-2 compared to Tcm (Fig. 2 *D*). Next, given the robust expression of effector molecules in terminal-Tem (Fig. 1 *E*), we evaluated granzyme abundance and the effector capacity of the distinct memory populations. Consistent with gene-expression levels, terminal-Tem had elevated levels of both GzA and GzB molecules compared to Tem and Tcm, while Tem had elevated expression of GzA and GzB compared to Tcm (Fig. 2 *E*). To determine how the diverse functional, phenotypic, and migratory/localization qualities of terminal-Tem are integrated to mediate pathogen clearance, we transferred sorted populations of congenically distinct terminal-Tem, Tem, and Tcm OT-I cells into naive recipient mice subsequently challenged with *Listeria monocytogenes* expressing OVA. We found that terminal-Tem conferred equivalent protection compared to Tem despite limited recall expansion (Fig. 2 *F*). Therefore, terminal-Tem have the most potent effector activity on a per cell basis following *Listeria* challenge, likely due to constitutively elevated expression of granzymes, perforin, and key migratory molecules (Fig. 1 *E*, 2 *E*). These data also demonstrate that the redefined population of Tem are functionally distinct from Tcm, which is notable as our reclassification suggests greater than previously appreciated similarity between Tem and Tcm, especially regarding homeostatic capacity and recall potential. The functional relevance of the three circulating memory populations was also investigated in a tumor vaccination model, in which cognate peptide and polyIC were administered following adoptive transfer of sorted terminal-Tem, Tem, and Tcm P14 cells into B16-GP_33-41_ tumor-bearing mice (*SI Appendix,* Fig. 3). Consistent with results from similar experiments (13, 54), Tcm conferred the greatest protection, likely due to their elevated lymph node homing capacity, recall proliferation, and enhanced survival. However, terminal-Tem provided minimal protection despite elevated expression levels of cytotoxic and migratory molecules, thus highlighting that the activity and protective role of each memory subset is likely dependent on the disease or therapeutic context.

**Figure 3.**
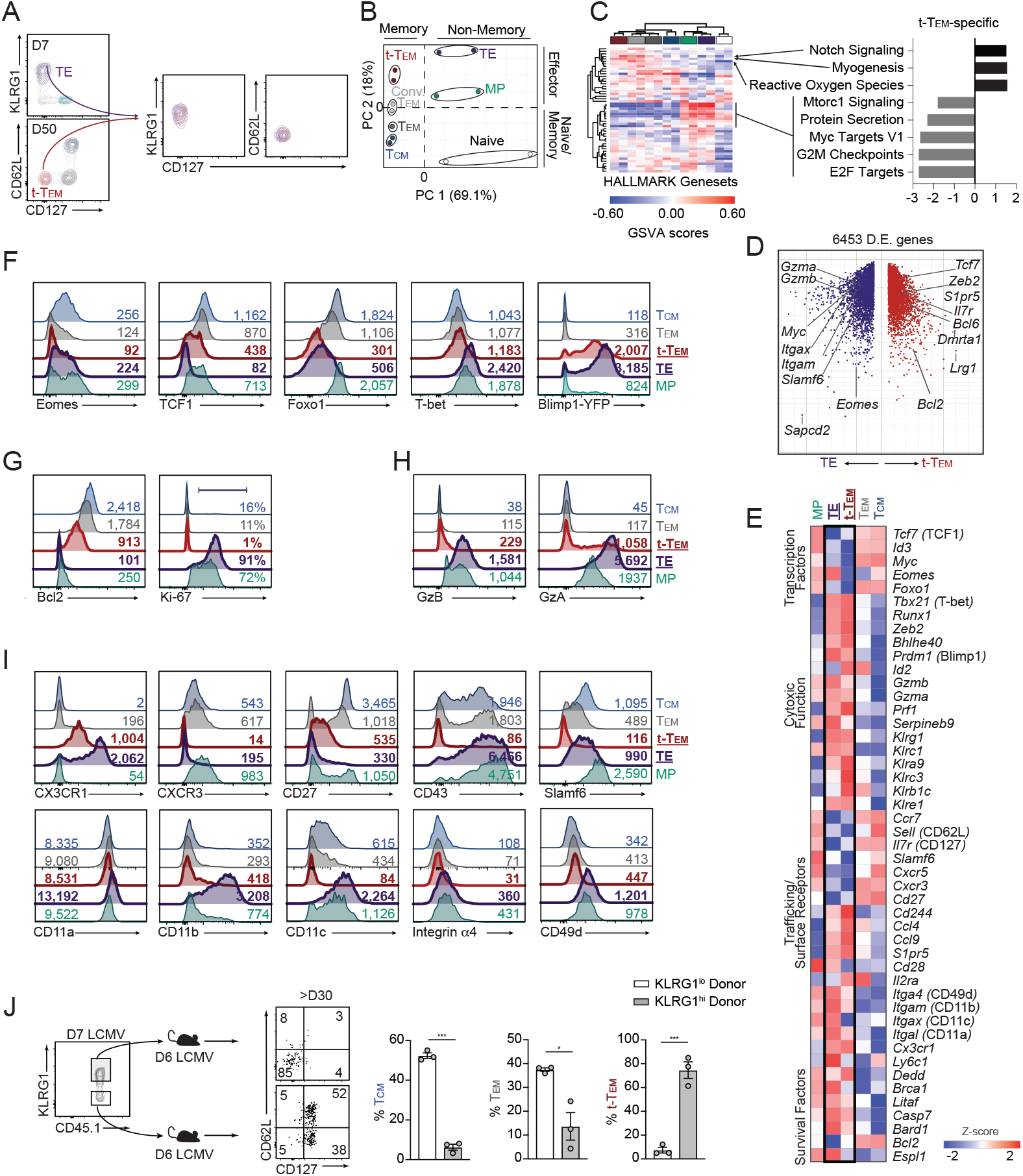
Terminal-Tem are memory T cells with features of terminal effector CD8 T cells. Memory cells were generated as in Figure 1. *(A)* Flow cytometric analyses indicating representative expression of KLRG1, CD62L, and CD127 by splenic P14 CD8 T cells on days 7 or 50 of infection. *(B)* Principal component analysis of RNAseq data comparing memory (Tcm, Tem, and terminal-Tem from day 55 of infection), effector (TE and MP from day 7 of infection), and naive P14 populations. *(C)* Gene set variation analysis (GSVA) of HALLMARK genesets, wherein rows and columns are ordered by hierarchical clustering (left) with key molecular processes unique to terminal-Tem compared with all other populations highlighted (right). Heatmap populations are denoted by color as in B. *(D)* Volcano plot demonstrating 6453 differentially expressed genes between TE (upregulated genes in purple) versus terminal-Tem P14 cells (upregulated genes in red), with key genes highlighted. *(E)* Heatmap of select genes differentially expressed by TE, MP, terminal-Tem, Tem, and Tcm. *(F-I)* Representative expression of transcription factors (F), homeostasis molecules *(G)*, granzymes *(H)*, and receptors/trafficking molecules *(I)* on TE (purple), MP (green), terminal-Tem (red), Tem (grey), and Tcm (blue) from spleens on day 7 (TE/MP) or 55 post LCMV infection. *(J)* KLRG1^hi^ or KLRG1^lo^ P14 CD8 T cells were sorted from spleens on day 7 of LCMV infection (right) and transferred into infection-matched, congenically distinct mice (left); Representative phenotype of donor cells and quantification of frequency of each memory population (right) >30 days after transfer. Numbers in plots represent the frequency of cells in the indicated gate or gMFI (F-J). All data are from one representative experiment out of 3 independent experiments *(A)* or 2 independent experiments *(F-J)* with n=3-4 per group; ***P<0.005. Graphs indicate mean ± s.e.m, symbols represent an individual mouse *(J)*.

### Terminal-Tem possess key characteristics of memory T cells with features of terminal effector CD8 T cells

The relatively low expression levels of CD62L and CD127 and elevated levels of KLRG1 on terminal-Tem resembled the surface phenotype of TE cells (Fig. 3 *A*), prompting the questions: 1) At late infection time points are terminal-Tem true memory cells, and 2) How are terminal-Tem, which phenotypically resemble short-lived TE cells, able to persist for >200 days following infection? Principal component analysis (Fig. 3 *B*) and a gene-expression similarity matrix (*SI Appendix,* Fig. 4) revealed that while terminal-Tem were transcriptionally distinct from D7 TE cells and more transcriptionally related to memory populations, terminal-Tem do exhibit some transcriptional similarly to TE relative to Tcm and Tem (Fig. 3 *B*, *SI Appendix,* Fig. 4). Further, we utilized HALLMARK genesets of key molecular processes for an unbiased evaluation of the biological relationship of terminal-Tem to memory, effector, and naive cells. Notably, the terminal-Tem subset clustered more closely to memory populations, further supporting the idea that terminal-Tem are memory T cells (Fig. 3 *B*). Additionally, a key defining characteristic distinguishing terminal-Tem from all other CD8 T cell populations was an apparent deficiency of certain biological processes related to proliferation such as ‘Myc Targets,’ ‘G2M checkpoint,’ and ‘E2F targets’ (Fig. 3 *B*), while oxidative stress and notch signaling may be distinctive features of terminal-Tem.

**Figure 4.**
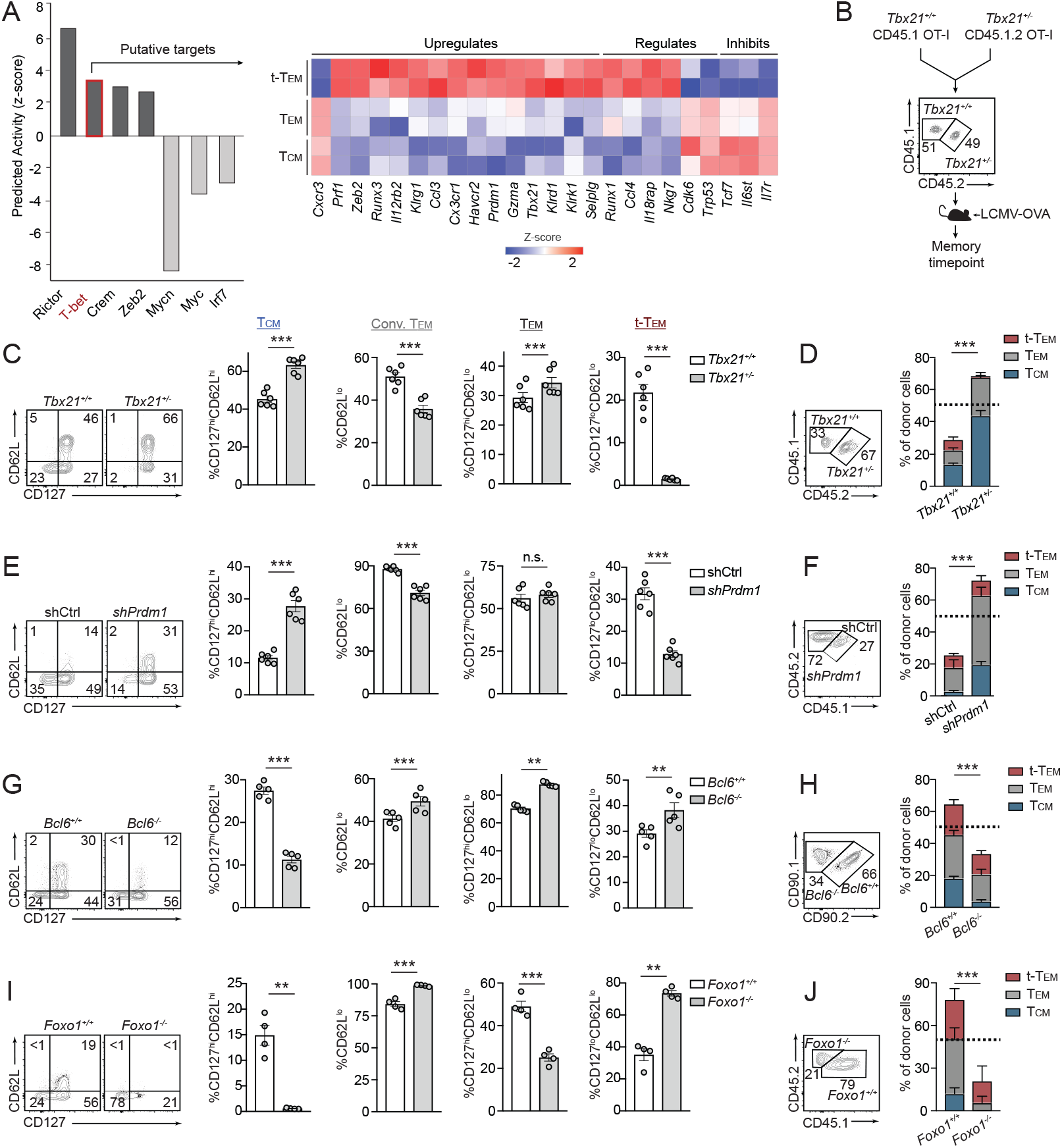
Terminal-Tem and Tem exhibit differential dependencies on key lineage-specifying transcription factors. *(A)* Regulatory transcription factors were predicted by Ingenuity Pathway Analysis based on gene-expression profiles of terminal-Tem, Tem and Tcm (left), and relative expression of key predicted targets of T-bet (right). *(B)* Summary of experimental model in which CD45.1 *Tbx21*^+/+^ OT-I cells were mixed 1:1 with CD45.1.2 *Tbx21*^+/−^ OT-I cells and transferred into recipient mice subsequently infected with LCMV-OVA. *(C)* Representative flow cytometry plot (left) and graphs of the frequency of splenic memory CD8 T cell subsets on day 72 of infection (right) from experimental schematic in *(B)*. *(D)* Representative flow cytometry plot indicating the ratio of recovered splenic *Tbx21*^+/+^ *and Tbx21*^+/−^ OT-I cells (left), and the relative abundance of terminal-Tem (red bars), Tem (gray bars), and Tcm (blue bars). *(E)* Congenically distinct P14 cells were transduced with *Prdm1* or *Cd19* (control) shRNA encoding retroviruses, mixed 1:1, and transferred into recipient mice subsequently infected with LCMV. Representative flow cytometry plot (left) and quantification of the frequency of splenic memory CD8 T cell subsets on day 25 of infection. *(F)* Representative flow cytometry plot indicating the ratio of recovered shCtrl *and shPrdm1* P14 cells (left), and the relative abundance of each subset (right). *(G)* Congenically distinct CD4-Cre-*Bcl6*^*fl/fl*^ P14 and *Bcl6*^*fl/fl*^ P14 cells were mixed 1:1 and transferred to recipient mice subsequently infected with LCMV. Representative flow cytometry plot (left) and quantification of the frequency of splenic memory CD8 T cell subsets on day 35 of infection (right). *(H)* Representative flow cytometry plot indicating the ratio of recovered *Bcl6*^*fl/fl*^ *and Bcl6*^−/−^ P14 cells (left), and the relative abundance of each subset (right). *(I)* Congenically distinct dLck-Cre-*Foxo1*^*fl/fl*^ P14 and *Foxo1*^*fl/fl*^ P14 cells were mixed 1:1 and transferred to recipient mice subsequently infected with LCMV. Representative flow cytometry plot (left) and quantification of the frequency of splenic memory CD8 T cell subsets on day 30 of infection (right). *(J)* Representative flow cytometry plot indicating the ratio of recovered *Foxo1*^*fl/fl*^ *and Foxo1*^−/−^ P14 cells (left), and the relative abundance of each subset (right). All data are from one representative experiment of 2 independent experiments with n=4-6 per group *(C-I)*; *P<0.05, **P<0.01, ***P<0.005. Graphs indicate mean ± s.e.m, and symbols represent an individual mouse.

Next, we sought to further understand how CD127^lo^CD62L^lo^KLRG1^hi^ (i.e. seemingly terminal or short-lived cells) CD8 T cells are able to persist for extended periods of time into the memory phase of infection. We compared the transcriptome and molecular profile of TE and terminal-Tem, which revealed a number of unique factors that may facilitate the surprising long-lived nature of terminal-Tem, including elevated levels of *Tcf7*/TCF1 and *Bcl2*/Bcl2 and lower levels of *Prdm1*/Blimp1 compared to TE (Fig. 3 *D-F*). However, terminal-Tem also shared a number of characteristics with TE cells relative to Tcm or Tem including expression of elevated levels of TE-signature molecules such as members of the killer cell lectin-like receptor family (e.g. *Klrg1*, *Klrc1, Klra9*), *Prdm1*/Blimp1, and effector molecules such as granzymes and perforin (Fig. 3 *D-H*). Further, consistent with the biological pathway analysis (Fig. 3 *C*) and Fig. 2 *C*, terminal-Tem exhibited minimal homeostatic proliferation evidenced by diminished Ki-67 staining whereas nearly all TE cells expressed Ki-67 (Fig. 3 *G*). Taken together, terminal-Tem are more “effector-like” than Tem and Tcm but also acquire key memory characteristics compared to TE, which may allow them to persist to memory timepoints.

To understand the ontogeny of terminal-Tem, we adoptively transferred sorted KLRG1^hi^ and KLRG1^lo^ populations from day 7 infected mice into infection-matched recipient mice; approximately 30 days post-infection, we evaluated memory T cell populations (Fig. 3 *J*). Terminal-Tem were primarily derived from KLRG1^hi^ cells, whereas Tem and Tcm were primarily derived from KLRG1^lo^ cells. Thus, terminal-Tem and Tem have unexpectedly distinct developmental pathways. These studies further clarify the ontogeny of memory T cells, demonstrating that KLRG1^lo^ effector cells are indeed the precursors of longer-lived memory cells (i.e. Tem and Tcm) but are not necessarily the originating population of all memory T cells, consistent with the finding that certain ex-KLRG1 cells can give rise to memory populations (13). Without parsing terminal-Tem from the Tem population, it would potentially be inferred that Tem are derived from both KLRG1^hi^ and KLRG1^lo^ effector cells.

### Terminal-Tem and Tem exhibit differential dependencies on key lineage-specifying transcription factors

We next utilized Ingenuity Pathway Analysis software to predict transcription factors putatively supporting terminal-Tem relative to Tem and Tcm lineages (Fig. 4 *A*). Rictor, T-bet, and Zeb2 were among the top predicted candidates promoting the terminal-Tem fate; all three factors have previously established roles in supporting TE formation (11, 17, 18, 55). T-bet is also widely considered to be essential for conventional Tem formation (3, 56); however, the computational approach predicted T-bet to be more influential in supporting terminal-Tem compared to both Tem and Tcm. To evaluate the fate-specifying role of T-bet on Tem and terminal-Tem differentiation, we co-transferred *Tbx21*^+/+^ or *Tbx21*^+/−^ OT-I cells at 1:1 ratio into congenically distinct recipient mice subsequently infected with LCMV-OVA (Fig. 4 *B*). This mixed transfer approach permitted investigation of a cell-intrinsic role for T-bet in regulating memory subset formation. Consistent with current paradigms, *Tbx21* heterozygosity resulted in a reduced frequency of conventional Tem (3, 56) (Fig. 4 *C*); however, the frequency of the redefined CD127^hi^CD62L^lo^ Tem population was unexpectedly unchanged or even slightly enhanced by diminished T-bet levels (Fig. 4 *C*). Notably, there was a dramatic loss in the frequency of terminal-Tem (Fig. 4 *C*), concurrent with the computational predictions (Fig. 4 *A*). On closer examination, the population frequencies did not necessarily translate to absolute abundance, and a key advantage of the co-transfer system is that it provides a sensitive method for directly comparing the relative abundance of *Tbx21*^+/+^ or *Tbx21*^+/−^ memory populations given the input ratio of transferred cells was 1:1. With this analysis, we found that *Tbx21* heterozygosity resulted in enhanced accumulation of total splenic OT-I cells at a memory timepoint compared to *Tbx21*^+/+^ OT-I cells (Fig. 4 *D*), and notably, *Tbx21* heterozygosity resulted in >2-fold increase in Tem (gray bars) but a nearly complete loss of terminal-Tem (red bars, Fig. 4 *D*). In summary, clarification of memory T cell nomenclature provided a framework for predicting critical roles for key regulatory molecules (e.g. T-bet, Zeb2, and Rictor) and allowed fine-tuning of the lineage-specifying role of T-bet, ultimately revealing that T-bet is essential for terminal-Tem but actually suppresses the formation of Tem and Tcm.

A distinguishing characteristic of terminal-Tem included elevated expression levels of Blimp1 compared to Tem and Tcm (Fig. 3 *F*). Similar to T-bet, Blimp1 is widely considered to be critical for Tem formation (3, 56), and given our relatively unexpected finding that T-bet was not essential for Tem formation, we next evaluated a role for Blimp1 in differentially supporting terminal-Tem and Tem populations. *Prdm1* RNAi resulted in a reduced percentage of CD62L^lo^ conventional Tem (Fig. 4 *E*), consistent with current paradigms (3, 56). However, we found that Blimp1-deficiency did not alter the frequency of the redefined Tem population (Fig. 4 *E*). Analogous to diminished T-bet levels, knockdown of *Prdm1* resulted in a greater accumulation of total P14 cells in the spleen compared to control P14 cells at a memory timepoint (Fig. 4 *F*), and consistent with previous reports (15, 16), Blimp1-deficiency resulted in a greater abundance of Tcm (blue bars, Fig. 4 *F*). However, despite robust *Prdm1* expression in the terminal-Tem population, *Prdm1* knockdown did not impair the overall formation of terminal-Tem (red bars) and actually enhanced the accumulation of Tem (grey bars) in the spleen (Fig. 4 *F*). Therefore, we also clarify the regulatory role of Blimp1 as a suppressor of Tem and Tcm formation.

We next examined the memory lineage-specifying roles of Bcl6 and Foxo1, transcription factors canonically recognized to support Tcm formation relative to Tem (3, 25–29). As expected, loss of *Bcl6* or *Foxo1* resulted in a reduced frequency of Tcm (3, 25–28) (Fig. 4 *G, I*). However, *Bcl6*-deficiency resulted in the enhanced frequency of Tem and terminal-Tem, whereas *Foxo1*-deficiency resulted in a lower frequency of Tem but increased frequency of terminal-Tem. Converse to the effects of diminished expression of T-bet or Blimp1, the relative abundance of donor cells revealed conditional deletion of *Bcl6* or *Foxo1* resulted in a loss of total memory P14 cells, including fewer Tcm (blue bars) compared to control P14 cells (Fig. 4 *H, J*). Unexpectedly, we also found that loss of *Foxo1* also resulted in a reduced abundance of CD127^hi^CD62L^lo^ Tem (Fig. 4 *H, J*). Therefore, clarification of memory T cell nomenclature has implications for how the roles of critical fate-specifying transcription factors are understood. In summary, we established T-bet as a central regulator of terminal-Tem, demonstrated that Foxo1 was unexpectedly essential for optimal Tem formation, and found that T-bet and Blimp1 may actually suppress development of Tem.

### Molecular profile of human terminal-Tem revealed by single-cell RNAseq

Although molecules such as CD28, CD27, and CD43 have been used to subset human CD8 T cells (46), circulating memory T cell populations in humans are most often distinguished based on expression of CD45RO (or CD45RA^lo^) and CCR7 expression, wherein CD45RO^hi^CCR7^hi^ cells are considered Tcm and CD45RO^hi^CCR7^lo^ are Tem (3, 5, 31). Utilizing this paradigm for subsetting human memory T cells, we found that conventionally defined CD45RO^hi^CCR7^lo^ Tem exhibit heterogeneous expression levels of CD127 and CD27 (Fig. 5 *A*), analogous to conventionally defined mouse Tem (Fig. 1 *A*). Further, multiparameter mass cytometry (CyTOF) analysis of human PBMCs from 6 healthy donors also demonstrated heterogeneous expression of effector molecules perforin, granzyme B, and granulysin within the CCR7^lo^ memory CD8 T cell population (Fig. 5 *B*, *SI Appendix,* Fig. 5 *A, B*), supporting the presence of a functionally distinct human terminal-Tem population that would otherwise be grouped into conventionally defined Tem.

**Figure 5.**
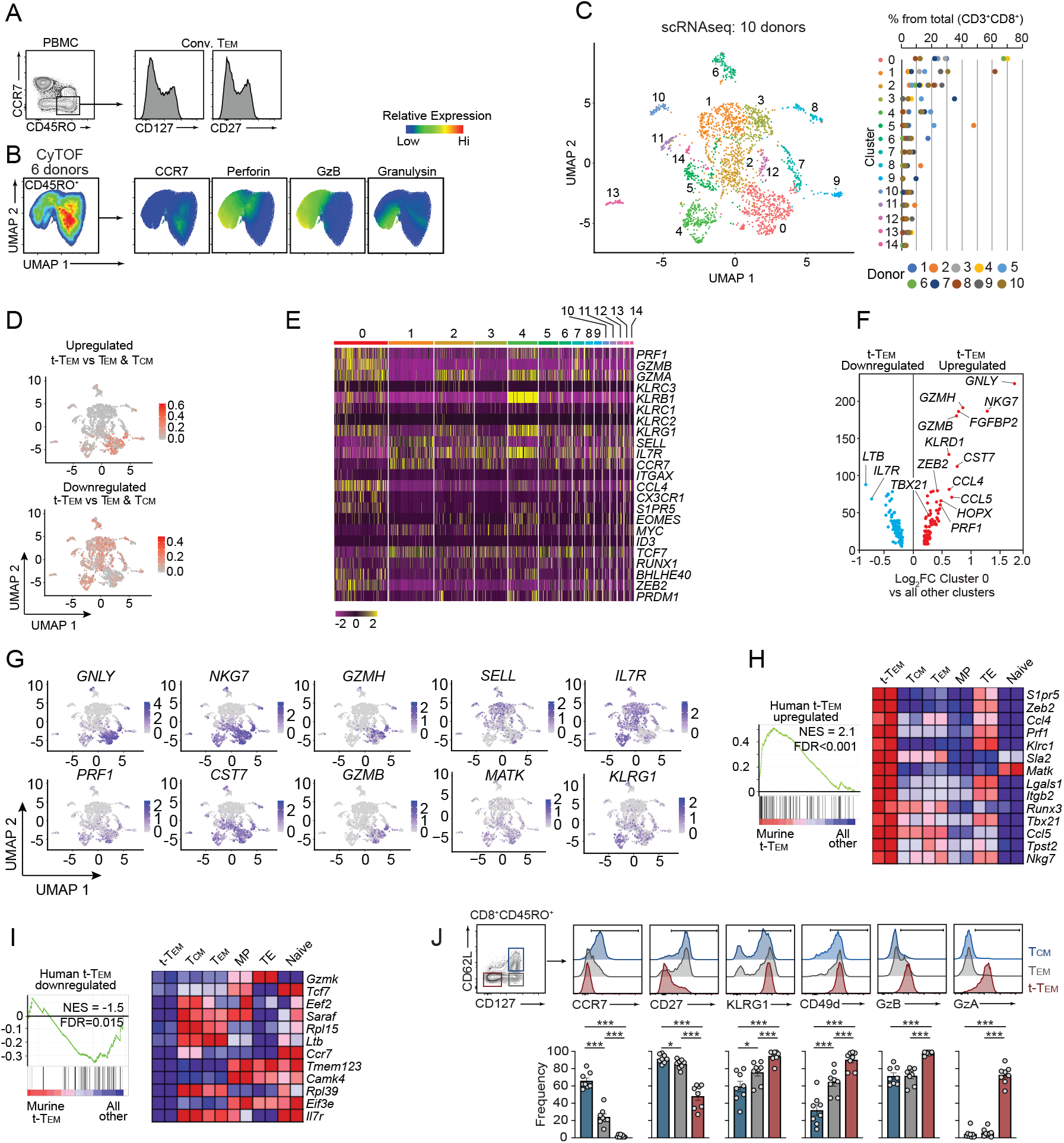
Molecular phenotype of human terminal-Tem. *(A)* Representative expression levels of CD127 and CD27 in conventional Tem (CD8^+^CD45RO^+^CCR7^−^) by flow cytometric analysis of PBMCs isolated from eight healthy donors. *(B)* PBMCs isolated from 6 healthy donors were analyzed by mass cytometry. Data was combined from all 6 donors, down-sampled to 3×10^4^ events, and UMAP highlighting heterogeneity of pre-gated CD45^+^CD3^+^CD8β+CD45RO^+^ T cells was constructed (left). Expression levels of CCR7, GzB, Perforin, and Granulysin are highlighted (right). *(C-I)* PBMCs isolated from 10 healthy donors, were processed for single-cell RNAseq analysis utilizing the 10x genomics platform. *(C)* CD8 T cells were pre-filtered based on expression of *CD3D*, *CD3E*, *CD8A*, and *CD8B*, and unbiased UMAP analysis revealed 15 distinct clusters (left). Frequency of each cluster per donor (right); each donor is designated by a distinct color indicated below. *(D)* Relative enrichment of the murine-derived terminal-Tem gene-expression signature (compared to both Tem and Tcm, see Supplemental Figure 6B) was evaluated within the human CD8 T single cell clusters *(E)* Relative expression of key molecules between the 15 distinct clusters from *(C)*. *(F)* Differential expression of highlighted genes uniquely upregulated or downregulated in cluster 0 (putative terminal-Tem population) compared with all other clusters. *(G)* Relative expression of indicated genes from *(C)*. *(H)* Gene set enrichment analysis of the human terminal-Tem gene-expression signature (genes uniquely upregulated in cluster 0 compared to all clusters) in murine terminal-Tem compared with all other murine populations (i.e. Tem, Tcm, TE, MP, and naive P14 cells), and (right) relative expression in murine populations of leading edge genes upregulated in both murine and human terminal-Tem. *(I)* Gene set enrichment analysis of the human non-terminal-Tem gene-expression signature (genes downregulated in cluster 0 compared to all clusters) in murine terminal-Tem compared with all other murine populations (i.e. Tem, Tcm, TE, MP, and naive P14 cells), and relative expression (right) of key genes in murine populations of leading edge genes downregulated in both murine and human terminal-Tem (bottom). *(J)* Representative gating of human terminal-Tem, Tem, and Tcm populations based on expression of CD62L and CD127 (left). Mean expression levels of CCR7, CD27, KLRG1, CD49d, GzA, and GzB for each subset among 8 healthy donors (right). All data are from one representative experiment of 8 donors *(A & J),* 6 donors *(B),* 10 donors *(C-I).* *P<0.05, **P<0.01, ***P<0.005. Graphs indicate mean ± s.e.m, and symbols represent an individual donor.

As there is often a disconnect between ‘markers’ used to distinguish murine memory populations and human populations, we utilized single-cell RNA-seq analysis for unbiased identification of human terminal-Tem. We performed single-cell RNAseq analysis on PBMCs from 10 healthy individuals (Fig. 5 *C*, *SI Appendix,* Fig. 6 *A*) as well as utilized a publicly available human PBMC dataset (*SI Appendix,* Fig. 7 *A-D*). Through these analyses, we detected extensive heterogeneity and diverse CD8 T cell states ranging from naive to memory populations yielding a total of 15 discrete sub-populations (Fig. 5 *C*). We identified cluster ‘0’ as enriched with the murine terminal-Tem gene-signature as well as partial enrichment in clusters ‘2,’ ‘4,’ and ‘7’ (Fig. 5 *D*, *SI Appendix,* Fig. 6 *B*). Gene-expression analysis of key regulatory molecules also supported the finding that cluster ‘0’ likely represents a terminal-Tem population as this cluster exhibited upregulation of cytotoxic molecules (*GZMA, GZMB, PRF1*), members of the killer cell lectin-like receptor family (*KLRCC3, KLRB1, KLRC1, KLRC2, KLRG1*), *CX3CR1, CCL4, S1PR5, BLHE40, ZEB2,* and *PRDM1* as well as relatively low levels of *SELL, IL7R, CCR7, MYC, ID3*, and *TCF7* (Fig. 5 *E-G*). Further, through gene-set enrichment analyses we confirmed that the cluster ‘0’ gene-expression signature (i.e. genes uniquely upregulated in cluster ‘0’ compared to all other clusters) was enriched in murine terminal-Tem (compared to MP, TE, Tcm, Tem and naive cells) (Fig. 5 *H*) and a gene-expression signature of transcripts uniquely downregulated in human terminal-Tem was depleted from murine terminal-Tem compared to other memory, effector, and naive populations (Fig. 5 *I*). Last, in analyses of PBMCs from 8 healthy donors we found that subsetting human memory (CD45RO^+^) CD8 T cells based on CD127 and CD62L expression distinguished a human terminal-Tem population, and this approach more clearly delineated human memory populations compared to CCR7 expression levels alone (Fig. 5 *J*). These data were consistent with profiling of murine terminal-Tem in that our analyses of human CD8 T cells revealed terminal-Tem to be transcriptionally and phenotypically distinct from Tem and Tcm populations.

## Discussion

Defining memory CD8 T cell functional and phenotypic states is valuable for understanding the secondary immune response and the molecular regulation of memory T cell differentiation. For example, the detection of a given subset can be informative for predicting memory T cell protection in the context of a given pathogen or cancer (4, 42, 48, 54, 56–58). As previously recognized by others (16, 41, 47, 48), we noted heterogeneity within CD62L^lo^ memory T cells, including a population of CD127^lo^CD62L^lo^ cells able to persist for several months after infection; this finding prompted the investigation and clarification of the CD62L^lo^ memory T cell compartment. Here, we refine the definition of conventional CD62L^lo^ Tem to include expression of CD127 (16, 44, 47, 48, 59), ultimately allowing parsing and characterization of a CD127^lo^CD62L^lo^ terminally-differentiated Tem memory population (i.e. terminal-Tem) separate from Tem. A key finding from this study, distinct from prior investigations of long-lived or persisting CD8^+^ T cells (41–44), was that our revised framework for examining circulating memory cells permitted an improved understanding of Tem biology as well as clarification of the roles of widely studied lineage-specifying transcription factors.

Human memory T cells with discrete lymphoid homing properties were first described by Sallusto *et al*. (5), yielding the definition of canonical memory T cell subsets Tcm and Tem. While the ontogeny and defining attributes of Tcm and Tem have been somewhat contentious (2–4, 60), general features of Tcm include heightened presence in secondary lymphoid organs, greater recall potential, enhanced IL-2 production, and greater homeostatic fitness (proliferation and survival) (2, 56, 61). Our studies confirm previous findings describing Tcm but redefine the attributes of the Tem population. Conventional Tem are characteristically considered to be more cytolytic, have a greater capacity to traffic to infected sites, have limited recall proliferation potential, generally accepted to be more terminally fated, and do not persist to the same extent as Tcm (2, 4, 56, 61). Through our refining of Tem identity, we reveal that Tem exhibit an enhanced presence in lymphoid tissues, more robust IL-2 production and recall potential, greater than expected homeostatic fitness (i.e. elevated homeostatic proliferation and expression of survival molecules such as Bcl2), heightened multipotency with the ability to give rise to Tcm, and a distinct molecular phenotype compared to prior paradigms (2, 3). These discrete functional attributes make certain memory populations better suited for handling specific infections, wherein previous studies have demonstrated Tcm are superiorly equipped to provide protection against systemic LCMV Cl13 infection (4, 33) and malignancy (54), whereas conventional Tem confer enhanced protection to vaccinia virus infection (48) and in some cases *Listeria* infection (53). We found that terminal-Tem and redefined CD127^hi^CD62L^lo^ Tem conferred equivalent protection to *Listeria* infection, but terminal-Tem conferred the most robust protection on a per cell basis. Although terminal-Tem displayed higher expression of cytolytic genes and upregulation of granzymes, they also exhibited limited recall proliferation, decreased lymphoid tissue presence, and impaired cytokine production compared to Tem. Conversely, Tcm were most protective in a tumor vaccine model compared to both Tem and terminal-Tem, likely due to an enhanced lymph node homing capacity (resulting in increased access to vaccine antigen (54)) and heightened recall proliferation. Taken together, the relative protection afforded by each memory T cell subset is dependent on the context of the disease or therapy, and the contribution of each subset is a culmination of unique molecular features impacting their location, expansion or survival, and functional activity. Resolving the terminal-Tem population provides clarity to this notion, and may impact future immunotherapy approaches. In connection, we have also identified putative ‘markers’ of human terminal-Tem that could inform vaccination studies or provide insight for pinpointing therapeutic or pathogenic roles for each memory state in a given therapy or disease setting.

Through profiling of terminal-Tem gene-expression patterns, we demonstrated that terminal-Tem were transcriptionally distinct from Tem, Tcm, and Trm. Further, despite a phenotypic resemblance to TE cells, terminal-Tem were more transcriptionally related to memory cells and sort-retransfer experiments demonstrated terminal-Tem are able to persist for extended periods of time (although not to the same extent as Tem and Tcm). Therefore, we conclude that terminal-Tem are a genuine memory T cell population with robust features of TE cells. We also found that compared to TE, terminal-Tem exhibited upregulation of pro-survival molecule Bcl2 as well as pro-memory transcription factors TCF1 and Bcl6, providing a possible explanation of their prolonged survival and potentially facilitating a dual terminal effector/memory phenotype. In connection, we refine paradigms of transcription factor-mediated regulation of memory T cell fate. To date, T-bet (11), Blimp1 (15, 16), Zeb2 (17, 18), Stat4 (19), and Id2 (20–23) have been considered essential for the formation of TE or conventional Tem cells, whereas Eomes (24), Bcl6 (25, 26), Foxo1 (27–29), Stat3 (26), and Id3 (22, 30) have been linked to the differentiation of MP or Tcm (3). Here, we demonstrated that optimal formation of the redefined Tem population was not impaired by diminished T-bet or Blimp1 expression. Further, we also highlighted that Tem actually require the pro-memory transcription factors Bcl6 and Foxo1. Notably, we found that loss of T-bet, Bcl6, and Foxo1 resulted in a reduced abundance of terminal-Tem.

The physiological role of memory T cells has been linked to both the resolution as well as the instigation of disease (1, 3, 14, 62, 63). The overall protective or pathogenic impact of memory T cells for a given disease setting is determined by both the quantity and quality of responses, which is likely dependent on the overall makeup and phenotype of the memory T cell compartment. While, it is well established that the circulating memory CD8 T cell population displays meaningful heterogeneity, it is also clear that T cells can exist in a broad spectrum of cell states rather than discrete subsets (1), complicating categorization of memory T cells into distinct lineages. Nonetheless, defining distinct states and delineating characteristics of terminal-Tem, Tem, and Tcm emphasizes the division of labor within memory T cell populations and highlights how the respective attributes of each subset may be complimentary both temporally and anatomically. Clarification of memory T cell nomenclature and lineage-specific characteristics provides insight for understanding the complex activity and molecular regulation of memory cell states and holds implications for the timing and delivery of immunotherapies.

## Funding

This study was supported by NIH P01AI13212 (A.W.G. and J.T.C), NIH U19AI109976 (A.W.G.), the Kenneth Rainin Foundation (J.T.C), and NIH K99/R00 CA234430 (J.J.M.).

## Author contributions

J.J.M. and H.N. designed and performed experiments, analyzed the data, and wrote the manuscript; K.O. assisted with designing experiments, tissue processing, analysis; M.C.R. assisted with computational analyses and the single-cell RNA-seq analysis; C.T. assisted with designing experiments, tissue processing, analysis; A.D.D. assisted with mouse breeding, tissue processing, analysis. B.S.B. assisted with collection and analysis of human single-cell RNA-seq analysis. S.M.H. provided reagents and advice for the design of experiments; J.T.C. supervised the collection of human PBMC and subsequent single-cell RNAseq analyses; A.W.G. and J.J.M. supervised the project, designed experiments, and wrote the manuscript.

## Competing interests

A.W.G. serves on the Scientific Advisory Boards of Pandion Therapeutics and Arsenal Bio.

## Accession Numbers

Accession numbers will be provided prior to publication.

---

**Figure S1.**
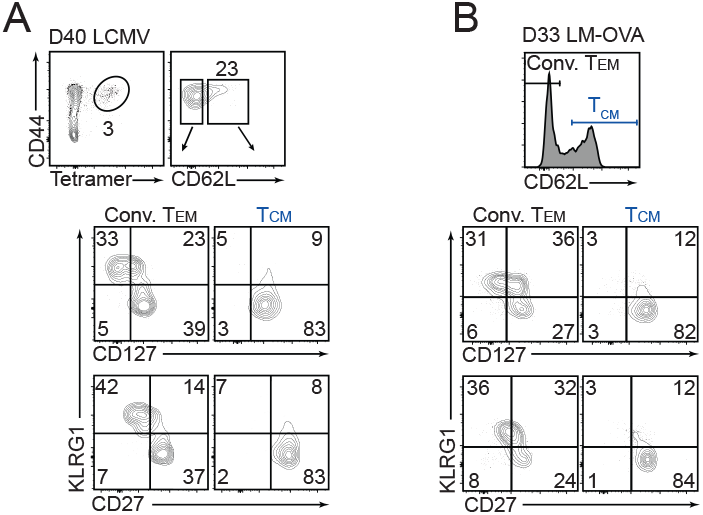
Heterogeneity within conventional Tem during acute LCMV infection and *Listeria monocytogenes* infection. *(A)* Representative expression of KLRG1, CD127, and CD27 by Tcm (CD62L^hi^) and conventional (Conv.) Tem (CD62L^lo^) on GP_33-41_-tetramer^+^ in PBL on day 40 of LCMV infection. *(B)* OT-I cells were transferred into congenically distinct recipient mice subsequently infected with *Listeria monocytogenes* expressing OVA (LM-OVA). Representative expression of KLRG1, CD127, and CD27 by Tcm (CD62L^hi^) and conventional (Conv.) Tem (CD62L^lo^) on OT-I cells in PBL on day 33 of LM-OVA infection. All data are from one representative experiment of 2 independent experiments with n=3-4 per group.

**Figure S2.**
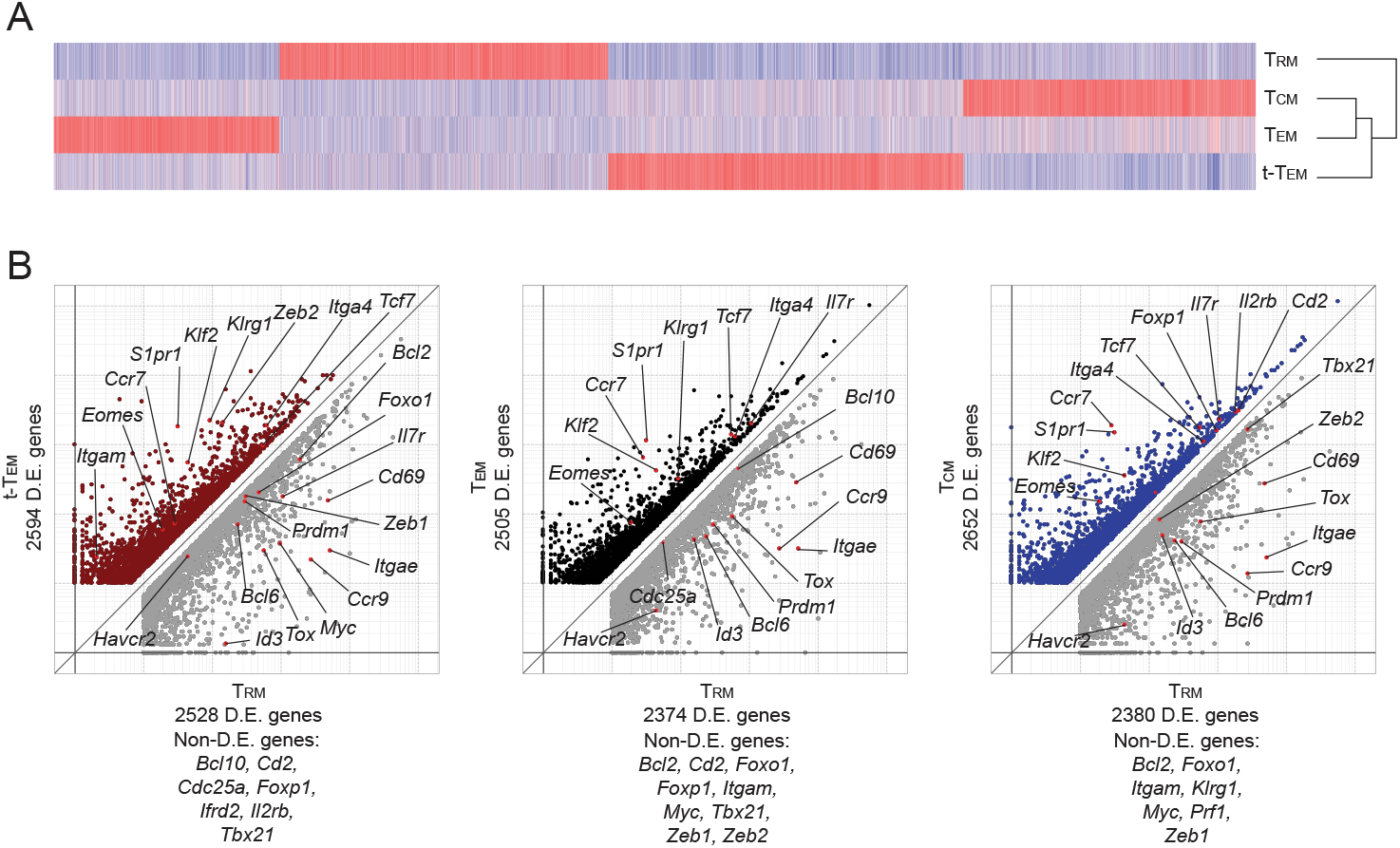
Transcriptional relationship of terminal-Tem, Tem, Tcm, and Trm. On day 55 of infection, splenic terminal-Tem, Tem, Tcm, and Trm P14 cells were sorted for RNA-seq analysis as described for Figure 1. *(A)* Heatmap illustrating differentially expressed genes (≥1.5-fold) ordered through hierarchical clustering between terminal-Tem, Tem, Tcm, and Trm. *(B)* Comparison of differential gene expression between Trm and each memory subset with highlighted genes differentially expressed or select non-differentially (non-D.E.) expressed genes listed below. All RNAseq samples consist of 2 biological replicates wherein each replicate is comprised of cells pooled from 2 mice.

**Figure S3.**
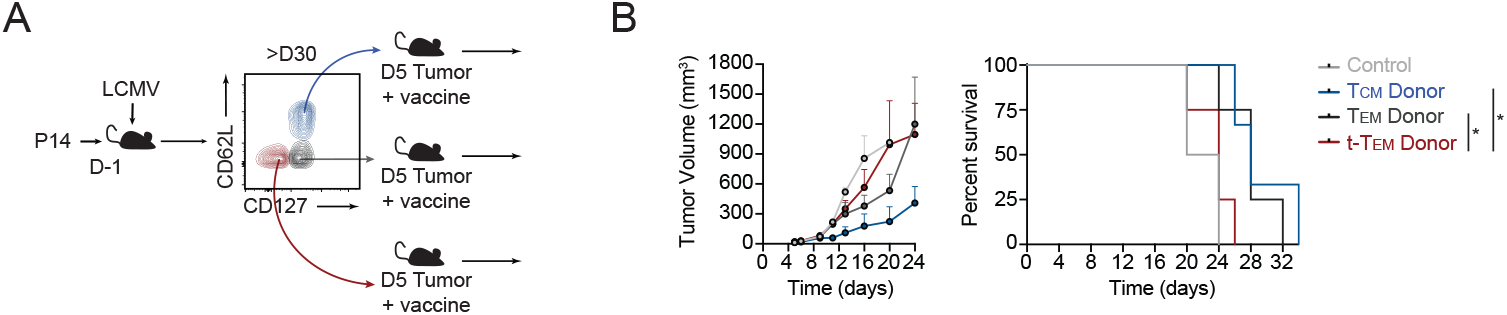
Terminal-Tem confer minimal protection in a tumor vaccine model. P14 CD8 T cells were transferred into congenically distinct mice that were infected with LCMV the following day. *(A)* Experimental schematic demonstrating at >30 days of infection, terminal-Tem (CD127^lo^CD62L^lo^), Tem (CD127^hi^CD62L^lo^), and Tcm (CD127^hi^CD62L^hi^) subsets were sorted and transferred to tumor-bearing mice. At the time of adoptive transfer, mice were immunized with 10 μg of GP_33-41_ and 2 μg of poly(I:C) in the ipsilateral flank. *(B)* Tumor volume (left) and survival curve of mice wherein mortality was defined as death, ulcerated tumor, or a tumor exceeding >1500mm^3^. Data are from one experiment with n=2 (control, no P14 transfer) or n=3-4 per group, *P<0.05. Graphs indicate mean ± s.e.m.

**Figure S4.**
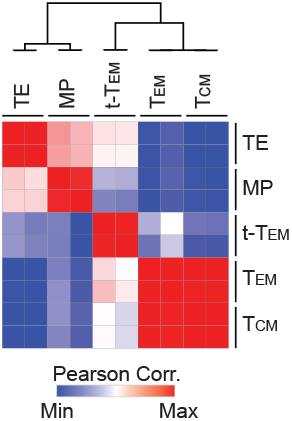
Gene-expression similarity matrix of terminal-Tem, Tem, Tcm, TE, and MP. Gene-expression similarity matrix (Pearson correlation) constructed based on differentially expressed genes between all populations of comparison from RNAseq data described in Figure 1.

**Figure S5.**
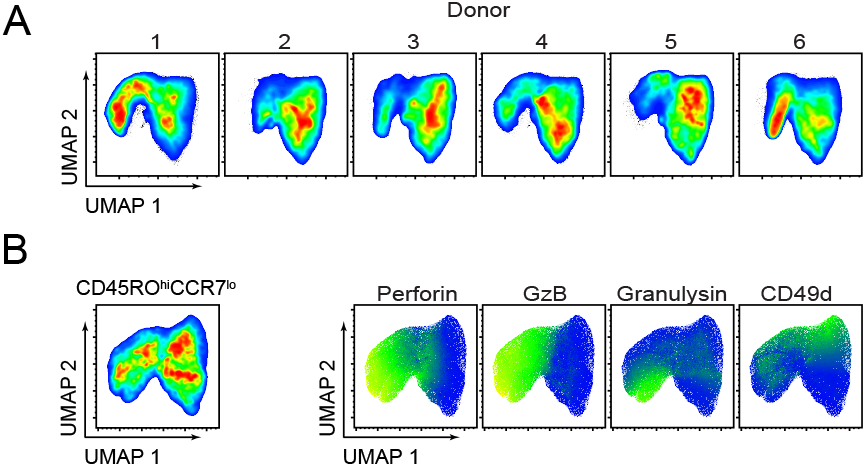
Mass cytometry analysis reveals heterogeneity of conventional human Tem. Mass cytometry was performed on human PBMCs from healthy donors (pertaining to Figure 5B). *(A)* UMAP of CD45^+^CD3^+^CD8β+CD45RO^+^ cells constructed for each of the 6 donors. *(B)* CD45^+^CD3^+^CD8b^+^CD45RO^+^CCR7^−^ (i.e conventional Tem) PBMCs from six donors integrated into one UMAP plot (left) and expression levels of highlighted molecules within this population (right).

**Figure S6.**
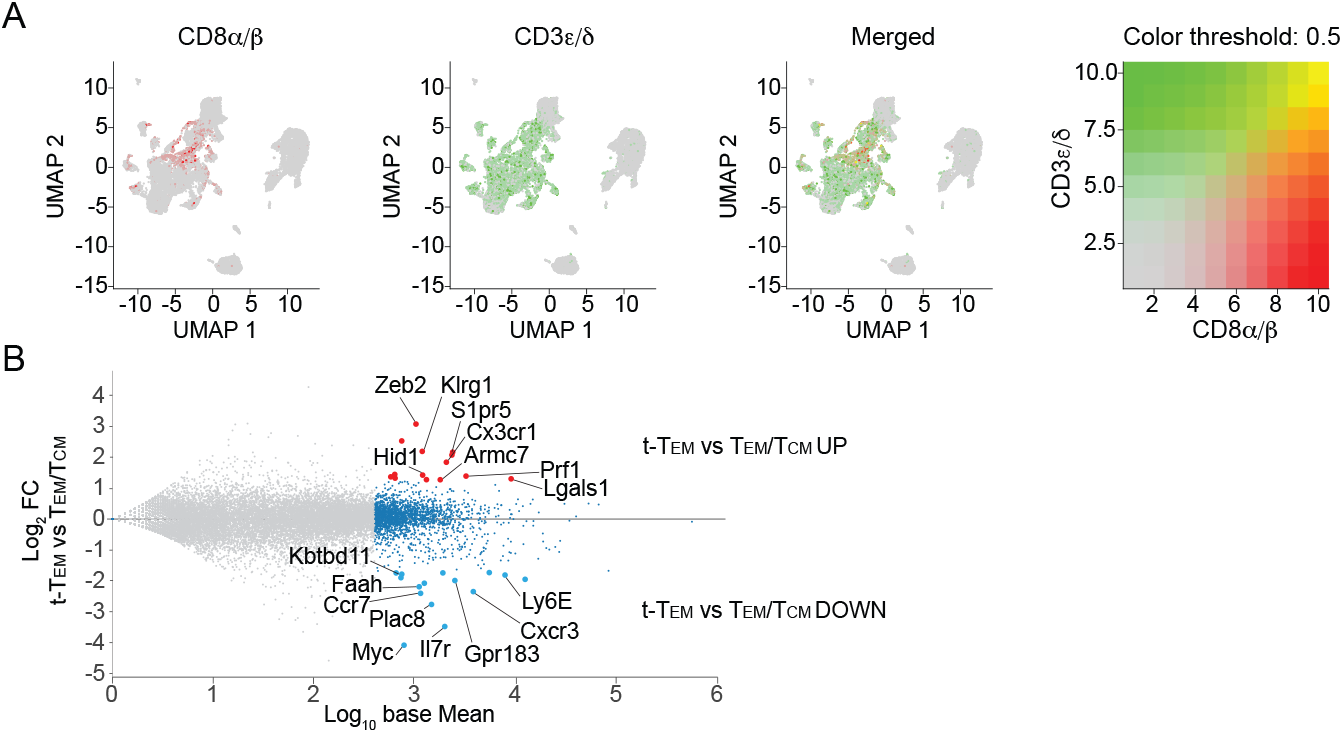
Identification of human terminal-Tem through single cell RNAseq analysis of human PBMCs. Single-cell RNAseq analysis was performed on human PBMCs from healthy donors (pertaining to Figure 5). *(A)* Expression patterns of *CD8A, CD8B, CD3D,* and *CD3E* in total PBMCs. CD8 T cells were filtered from total single-cell datasets based on expression of *CD8A, CD8B, CD3D,* and *CD3E* for subsequent analyses (Figure 5). *(B)* Establishment of gene signatures upregulated in murine terminal-Tem compared to both Tcm and Tem, as well as genes downregulated in terminal-Tem compared to Tcm and Tem. Gene sets were used for Figure 5*D*.

**Figure S7.**
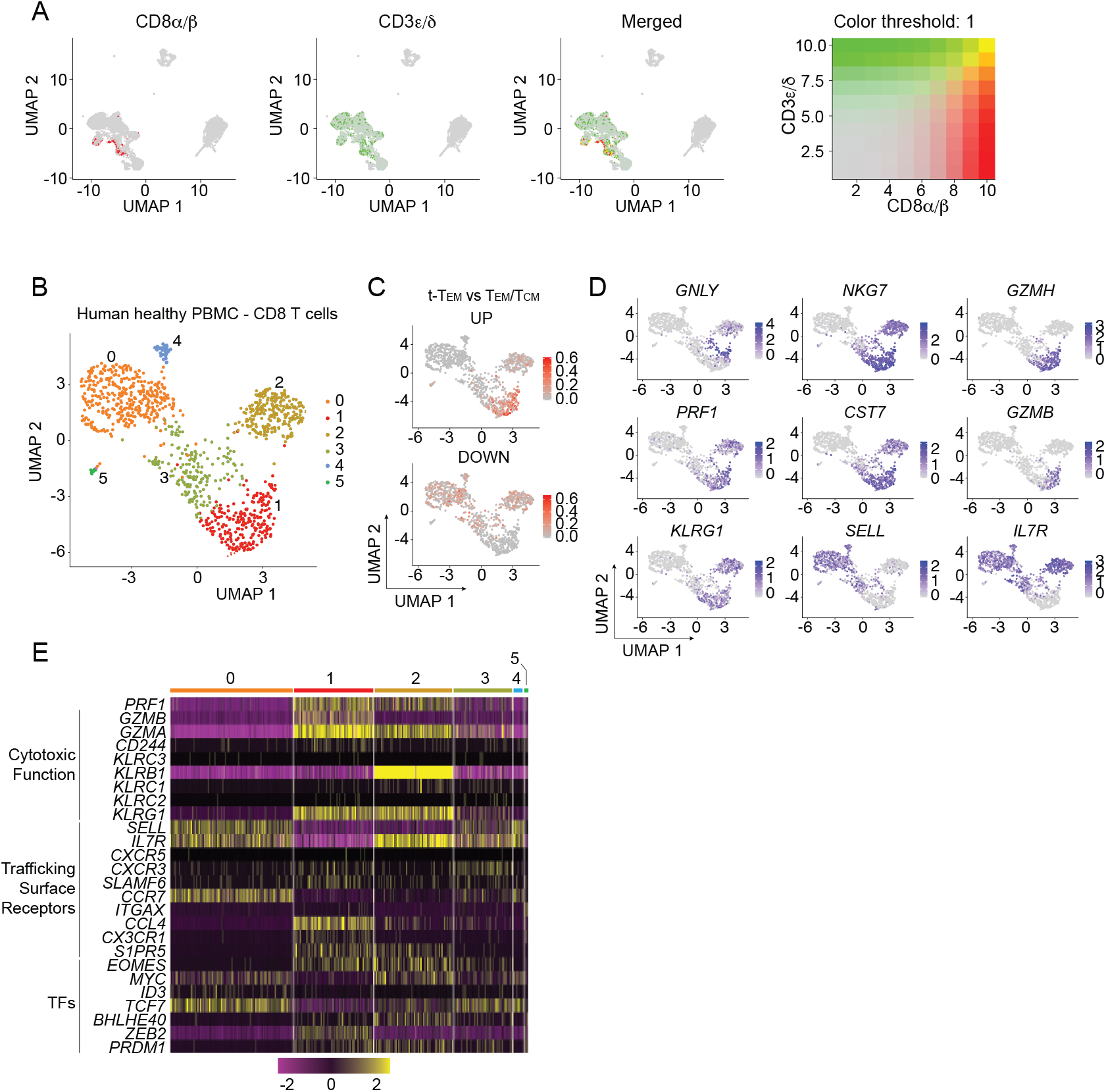
Phenotype of terminal-Tem from a publicly available human PBMC single-cell RNAseq dataset. The publicly available “10k PBMC scRNAseq” dataset from 10x Genomics was analyzed for identification of a human terminal-Tem population. *(A)* Expression patterns of *CD8A, CD8B, CD3D,* and *CD3E* in total PBMCs. CD8 T cells were filtered from all cells based on expression of *CD8A, CD8B, CD3D,* and *CD3E* for subsequent analyses. *(B)* Unbiased UMAP analysis revealed 6 distinct clusters of CD8 T cells *(C)* Relative enrichment of the murine terminal-Tem gene-expression signature was evaluated within the human CD8 T cells. *(D)* Relative expression of key molecules between the 6 distinct clusters from *(B)*. *(E)* Expression levels of highlighted genes between the 6 distinct clusters.

## Materials and Methods

### Mice

All mouse strains were bred and housed in pathogen-free conditions in accordance with the Institutional Animal Care and Use Guidelines of the University of California, San Diego. P14 mice (with transgenic expression of H-2D^b^-restricted TCR specific for LCMV glycoprotein 33-41, OT-I mice (with transgenic expression of H-2K^b^-restricted TCR specific for ovalbumin 257-264), *Tbx21*^+/−^ mice, *Bcl6*^*fl/fl*^*Cd4*^*Cre*^, *Bcl6*^*fl/fl*^*Ert2*^*Cre*^ mice, *Foxo1*^*fl/fl*^*dLck*^*Cre*^ mice, Blimp1-YFP reporter mice, CD45.1 mice, and Thy1.1, mice were purchased from Jackson Laboratory and bred at UCSD.

### Cell transfers and infections

To generate memory T cells, naive P14 or OT-I CD8 T cells (5×10^4^) were adoptively transferred into congenically distinct recipient mice, which were infected the next day with 2×10^5^ PFU LCMV-Armstrong i.p. or with 8×10^4^ CFU *Listeria monocytogenes-*OVA (LM-OVA) i.v. For KLRG1^hi^/KLRG1^lo^ transfers, P14 effector cells were sorted from spleens based on KLRG1 expression on day 7 of infection and 1×10^6^ cells were transferred into congenically distinct, infection matched mice. For fate studies, memory P14 cells were sorted >30 days post infection from spleens based on CD62L and CD127 expression, and 1×10^5^ cells were transferred into naive recipients.

For re-challenge assays, memory OT-I subsets were sorted from recipient spleen and lymph nodes at >30 days after LCMV-OVA infection and 1×10^5^ cells were transferred i.v. into congenically distinct naive recipients that were subsequently infected with 8×10^4^ CFU LM-OVA i.v. On day 3 of infection, spleens from recipient mice were homogenized in 0.2% igepal (Sigma-Aldrich) and serial dilutions in BHI media were plated onto BHI plates in the presence of antibiotic selection. Colonies were counted the following day and normalized to splenic weight. For tamoxifen-induced deletion, 1 mg of tamoxifen (Cayman Chemical Company) was emulsified in 100 μl of sunflower seed oil (Sigma-Aldrich) and administered i.p. from days 2-7 after infection.

To assess the activity of each memory subsets in a tumor vaccine model, 5×10^5^ B16-GP_33-41_ cells were transplanted s.c. into naive recipient mice and allowed to progress for 5 days. Next, 7.5×10^4^ congenically distinct cells of each sorted memory population were adoptively transferred and tumor-bearing mice were subsequently immunized with 10 μg of GP_33-41_ and 2 μg of poly(I:C) (GE Healthcare) s.c. into the ipsilateral flank. Tumors were monitored daily and mice with ulcerated tumors or tumors exceeding 1500mm^3^ were euthanized.

### Antibodies, flow cytometry and cell sorting

CD8 T cell subsets were first gated on congenically distinct P14 cells or CD44^+^Tetramer^+^ cells (as indicated), and then defined as: terminal-Tem: CD127^lo^CD62L^lo^; Tem: CD127^hi^CD62L^lo^; Tcm: CD127^hi^CD62L^hi^, conventional Tem: CD62L^hi^; TE: D7 KLRG1^hi^CD127^lo^; MP: D7 KLRG1^lo^CD127^hi^. The following antibodies were used for murine surface staining: CD8 (53-6.7), CD27 (LG-7F9), CD43 (1B11), CD44 (IM7), CD45.1 (A20-1.7), CD45.2 (104), CD62L (MEL-14), CD127 (A7R34), CX3CR1 (SA011F11), KLRG1 (2F1), CXCR3 (CXCR3-173), Slamf6 (13G3), CD11a (M17/4), CD11b (M1/70), CD11c (N418), CD49d (R1-2). Cells were stained for 20min at 4°C in PBS supplemented with 2% bovine serum albumin and 0.1% sodium azide. Intracellular staining was performed with the Foxp3-transcription factor staining buffer kit (eBioscience) using the following antibodies: IL-2 (JES6-5H4), IFNγ (XMG1.2), TNFα (MP6-XT22), Bcl2 (3F11), GzA (CB9), GzB (GB12), Ki-67 (SolA15), T-bet (4B10), Foxo1(C29H4), Eomes (Dan11mag), TCF1 (C63D9). For cytokine staining, memory populations sorted from spleens at >30 days after LCMV infection were incubated for 4 hours at 37°C with 10 nM GP_33-41_ peptide, congenically distinct naive splenocytes, and CD107a (1D4B) antibody was included in the media for the entirety of the stimulation; protein transport inhibitor was added after the first hour of incubation. For analysis of human PBMCs, the following antibodies were used: CD8 (SK1), CD45RO (UCHL1), CCR7 (G043H7), CD62L (DREG56), CD127 (A019D5), CD27 (M-T271), KLRG1 (13F12F2), CD49d (9F10), GzA (CB9), GzB (GB11). Flow cytometry data were generated with an LSR Fortessa or LSR Fortessa X-20 (BD) and FlowJo Software (TreeStar) was used for analyses. Cells were sorted using FACS Aria, FACS Aria Fusion, or Influx (BD) instruments.

### RNAi studies

shRNAmirs targeting mouse *Prdm1* or *Cd19* (control) cloned into a pLMPd-Amt vector were utilized, and retroviral supernatants were generated as previously described(32). For transductions, naive P14 cells from spleen and lymph nodes were enriched via negative selection using MACS magnetic beads according to the manufacturer’s protocol (Miltenyi Biotec). Enriched P14 cells (2×10^6^) were activated in 6-well plates coated with 100 μg/mL goat anti-hamster IgG (H+L, Thermoscientific) and 1 μg/mL anti-CD3 (145–2C11) and 1 μg/mL anti-CD28 (37.51) (eBioscience) for 18 h. Culture media was replaced with retroviral supernatant supplemented with of 50μM BME and 8 μg/mL polybrene (Millipore), and cells were centrifuged for 60min at 2000rpm, 37°C. After 24 h, P14 cells transduced with *Prdm1* shRNAmir encoding-or *Cd19* shRNAmir encoding-retroviruses were mixed at 1:1 ratio and a total of 5×10^5^ cells were transferred to congenically distinct recipient mice that were infected with LCMV-Armstrong.

### RNA-seq analysis

On day 55 of LCMV-Armstrong infection, 1×10^3^ splenic terminal-Tem (CD127^lo^CD62L^lo^), Tem (CD127^hi^CD62L^lo^), Tcm (CD127^hi^CD62L^hi^), conventional Tem (CD62L^lo^), and small intestine epithelial Trm (CD103^hi^CD62L^lo^) P14 cells were sorted into TCL buffer (Qiagen) with 1% 2-Mercaptoethanol. RNA was isolated and RNA-seq library preparation was carried out as per Immgen Protocols (https://www.immgen.org/Protocols/11Cells.pdf). D7 MP and TE as well as naive RNAseq datasets were previously published (32). Differentially expressed genes were identified through the Multiplot Studio module within Genepattern with >1.5 fold cutoff and expression threshold >10. Heatmaps were generated using Morpheus software (https://software.braodinstitute.org/morpheus). K-means clustering was set according to the number of groups, maximum number of iterations = 1000. Triwise plots as well as rose plots were created with the log_2_ transformed data using the Triwise R package and following https://zouter.github.io/triwise/rd.html. PCA was performed in R studio using Deseq2 normalized expression values. GSVA scores for each sample were calculated using GSVA R module (64) and the Hallmarks.v7 geneset collection (MSigDb, Broad Institute). Unsupervised clustering was performed with one minus Pearson correlation metric. GSEA was run with a Signal to noise ratio for metric ranking, 1000 permutations based on gene set and the Mouse Gene Symbol Remapping MSigDB.v7.0 chip. ‘Memory’ and ‘effector’ genesets were generated from analysis of populations from the Immgen database (www.immgen.org); specifically, D8 OT-I and D100 OT-I cells from LM-OVA mice were compared and ‘memory’ genes were defined as upregulated in the D100 population relative to the D8 population and ‘effector’ genes were downregulated in the D100 population relative to the D8 population. The ‘proliferation’ geneset was from module 2 of Best et al. (52). For computational prediction of putative regulators of terminal-Tem (Fig. 4a), the Inguenity Pathway Analysis (Qiagen, https://www.qiagenbioinformatics.com/products/ingenuity-pathway-analysis/) was used. Naive P14, LCMV D7 TE, LCMV D7 MP RNAseq datasets are from Milner et al. (32).

### Collection of human PBMCs

The Human Research Protection Programs at the University of California, San Diego and the San Diego VA Healthcare System approved the study. For mass cytometry, peripheral blood was obtained from patients undergoing colonoscopy at the University of California, San Diego and the San Diego VA Healthcare System after obtaining informed consent. Healthy individuals were undergoing colonoscopy as part of routine clinical care for colorectal cancer screening/surveillance or non-inflammatory gastrointestinal symptoms that included constipation or rectal bleeding. Inclusion criteria included age over 18 years old and absence of significant comorbidities or colorectal cancer. Blood was collected in BD Vacutainer CPT tubes and centrifuged at 400 x g for 25 minutes. The buffy coat layer was removed, washed, and counted. Cells were resuspended in freezing buffer (10% DMSO, 40% complete RPMI [RPMI +10% fetal bovine serum (FBS) +100 U/mL penicillin/100 μg/mL streptomycin], 50% FBS), placed into a freezing container, and stored at −80°C. Cells were recovered, washed, filtered, and labeled with anti-human CD45 mAbs. CD45^+^ immune cells were sorted on a FACSAria2 (BD Biosciences) or utilized for mass cytometry analysis. For flow cytometry analysis of intracellular and surface markers, buffy coats from healthy donors were purchased from Stanford Blood Center. PBMCs were isolated via Ficoll-Paque (GE Healthcare) density-gradient separation and flow cytometry analysis was performed immediately.

### Mass cytometry (CyTOF) of human peripheral blood mononuclear cells

Blood was pre-processed as above, washed with protein-free PBS (Rockland), and stained with Cell-ID cis-platin (Fluidigm) 5 minutes at room temperature, then washed with CyFACS buffer [PBS + 0.1%(w/v) bovine serum albumin (Sigma-Aldrich) + 2mM EDTA (Invitrogen) + 0.05%(v/v) NaN_3_)]. Staining and fixing cells for CyTOF using 29 metal-conjugated antibodies in accordance with cell staining protocols (Fluidigm). Cells were analyzed on a Helios CyTOF 2 mass cytometer (Fluidigm) at the La Jolla Immunology Institute Flow Cytometry Core at approximately 300 events/s. All antibodies used for mass cytometry were purchased from Fluidigm or conjugated using Maxpar Antibody Labeling Kit (Fluidigm) and are listed in table. FCS files created from mass cytometry were analyzed on FlowJo (BD Biosciences). UMAP (uniform manifold approximation and projection) for dimensional reduction was done on cells pre-gated on: Live cells, CD45^+^, CD3^+^, CD4^−^, CD8β+ cells, with down-sampling to 5×10^4^ cells per patient, and subsequent concatenation of all individuals (6 individuals in total). UMAP utilized protein expression of CCR7, CXCR5, Eomesodermin, Perforin, Granzyme B, Integrin α4 (CD49d), Granulysin, and NKG2A (KLRC1) for analysis.

### Single-cell RNAseq analysis

Cells were washed and resuspended in phosphate buffered saline (PBS) + 0.04% bovine serum albumin. Single cell libraries were prepared according to the protocol for 10X Genomics Chromium Single Cell Gene Expression. Approximately 2 ×10^4^ sorted CD45^+^ cells were loaded and partitioned into Gel Bead In-Emulsions (GEM). scRNA libraries were sequenced on a HiSeq4000 (Illumina). scRNAseq analysis was performed using cellranger software and the developmental version of Seurat 3.1 (65, 66) in R Studio. Cellranger was used with default parameters with the exception of cellranger aggr mapped=none. Seurat analysis of 10X count matrices was done by following these steps: low quality cells, identified by percent mitochondria<15, nFeature_RNA<200 or >2000, were removed, counts were normalized with SCTransform, dimensionality reduction and cluster identification were done with uMAP (dims=1:40), FindNeighbors (dims=1:40), and FindClusters (resolution=1). 10K PBMC scRNAseq dataset was downloaded from https://support.10xgenomics.com/single-cell-gene-expression/datasets/3.0.0/pbmc_10k_protein_v3. CD8 T cells were subsetted based on the expression of *CD3D, CD3E, CD8A* and *CD8B* expression. Differential expression of genes per cluster was done with FindAllMarkers function with default parameters and min.pct=0.1 and logfc.threshold=0.2.

### Statistics

Two-tailed, paired and unpaired Student’s *t* test was used for comparisons between two groups. Statistical analysis was performed using GraphPad Prism software. Log-rank (Mantel-Cox) test was used for survival curves. All statistical tests were performed with GraphPad Prism software. P<0.05 was considered statistically significant.

